# Coding of odour and space in the hemimetabolous insect *Periplaneta americana*

**DOI:** 10.1101/860908

**Authors:** Marco Paoli, Hiroshi Nishino, Einat Couzin-Fuchs, C. Giovanni Galizia

**Affiliations:** Department of Neuroscience, University of Konstanz, 78457 Konstanz, Germany; Research Institute for Electronic Science, Hokkaido University, Sapporo 060-0812, Japan

**Keywords:** olfaction, odour coding, calcium imaging, antennal lobe, insects, cockroach

## Abstract

The general architecture of the olfactory system is highly conserved from insects to humans, but neuroanatomical and physiological differences can be observed across species. The American cockroach, inhabiting dark shelters with a rather stable olfactory landscape, is equipped with long antennae used for sampling the surrounding air-space for orientation and navigation. The antennae’s exceptional length provides a wide spatial working range for odour detection; however, it is still largely unknown whether and how this is also used for mapping the structure of the olfactory environment. By selective labelling antennal lobe projection neurons with a calcium sensitive dye, we investigated the logic of olfactory coding in this hemimetabolous insect. We show that odour responses are stimulus-specific and concentration-dependent, and that structurally related odorants evoke physiologically similar responses. By using spatially confined stimuli, we show that proximal stimulations induce stronger and faster responses than distal ones. Spatially confined stimuli of the female pheromone periplanone-B activate sub-region of the male macroglomerulus. Thus, we report that the combinatorial logic of odour coding deduced from holometabolous insects applies also to this hemimetabolous species. Furthermore, a fast decrease in sensitivity along the antenna, not supported by a proportionate decrease in sensillar density, suggests a neural architecture that strongly emphasizes neuronal inputs from the proximal portion of the antenna.

**Summary statement:** By selective labelling the cockroach’s antennal lobe output neurons, we investigated the logic of olfactory coding in a hemimetabolous insect, showing that odour responses are stimulus-specific, concentration-dependent, and preserve information on the spatial structure of the stimulus.

## Introduction

The olfactory system is highly conserved across species (Galizia and Rössler, 2010; Hildebrand and Shepherd, 1997; Kay and Stopfer, 2006). Nevertheless, differences in the neural architecture and physiology can be observed. The American cockroach *Periplaneta americana* is a gregarious animal, often living in dark shelters with a rather stable olfactory landscape. It is equipped with exceptionally long antennae (up to 5 cm, often exceeding its body length) that actively sample the surrounding air-space for orientation and navigation. Long antennae allow a wide working range for odour detection. However, it is still largely unknown whether and how this could be used for mapping the spatial structure of the olfactory surrounding. Insects rely on bilateral comparison of antennal inputs and on their temporal integration for perceiving the distribution of odorants in space (Borst and Heisenberg, 1982; Gomez-Marin, 2010; Takasaki et al., 2012). Nonetheless, it has recently been suggested that odour source localization in cockroaches may rely more on the overall antennal length rather than on bilateral inputs (Lockey and Willis, 2015). Cockroach antennae are covered in sensilla, cuticular structures of different size and shape, together hosting approximately 220,000 olfactory sensory neurons (OSNs) (Sass, 1983; Schaller, 1978). Olfactory neurons originating in different locations along the antenna converge into the antennal nerve, and descend towards the antennal lobe (AL), the first odour processing centre, which is structured into ∼205 anatomical and functional units, named glomeruli (Watanabe et al., 2010). Each glomerulus receives inputs from on average 1,000 OSNs originating in different antennal segments (annuli), probably all expressing the same olfactory receptor gene. The male cockroach possesses an additional ∼36,000 OSNs for the detection of the minor and major components of the female sex pheromone, periplanone A and B, respectively (Sass, 1983; Schaller, 1978; Watanabe et al., 2012). Their axons innervate two exceptionally large glomeruli, which form the macroglomerular complex (Schaller, 1978; Watanabe et al., 2012). Glomeruli are interconnected by a dense network of local interneurons (Fusca et al., 2013; Husch et al., 2009), and processed olfactory information is relayed to higher order brain centres by uniglomerular (uPNs) and multiglomerular projection neurons (mPNs) (Malun et al., 1993). Uniglomerular PNs extend their dendritic arborization in a single glomerulus, thus receiving direct input from only one OSN type. Two subgroups of uPNs innervate distinct glomerular populations, their cell bodies clustering in two distinct groups in the antero-dorsal side of the AL (Strausfeld and Li, 1999; Watanabe et al., 2017). In the American cockroach, each ordinary glomerulus is innervated by only one uPN, whereas the macroglomerulus (MG) is innervated by multiple uPNs with a highly stereotyped topology (Galizia, 2018; Nishino et al., 2018). Two main PN axon bundles exit from the medio-dorsal side of the antennal lobe (note that all orientations here provided are in body axis system), and relay odour-related information to the protocerebrum. A first thick and more medial root splits in three distinct antennal lobe tracts (ALTs (Ito et al., 2014), previously termed antenno-cerebral tracts, ACTs (Malun et al., 1993)): ALT-I (also inner-ALT), ALT-II and ALT-III. They comprise the uPN axons, and innervate both the mushroom body (MB) calices and the lateral protocerebrum (LP). Posterior to the first root, a second bundle of axons exits the antennal lobe forming the ALT-IV and the outer-ALT. These lateral tracts collect the axons from mPNs, and innervate almost exclusively the lateral protocerebrum (Malun et al., 1993).

Cockroaches are hemimetabolous insects. Hence, the basic structures of the brain are established during embryogenesis, and gradually increase in volume during multiple post-embryonic moults. At each moult, new antennal segments are added proximally to the head, together with new OSNs (Schafer and Sanchez, 1973; Watanabe et al., 2018). As shown for the macroglomerulus (Nishino et al., 2018) and suggested for common glomeruli (Nishino and Mizunami, 2007), OSNs axons descend along the antennal nerve and terminate onto the antennal lobe glomeruli, forming segment (thus age)-specific layers of OSNs terminals from the core to the cortex of each glomerulus, producing an antennotopic innervation (Nishino et al., 2018). In this respect, they differ from holometabolous insects, which develop the complete glomerular organization during a single metamorphic event, and where the innervation topology does not reflect a sequence of post-embryonic moults (although it may still be antennotopic). In all insects, AL output neurons develop during embryogenesis. However, whereas the holometabolous insects studied so far have multiple uPNs per glomerulus (fruit fly: 50 glomeruli and 150-200 uPNs (Grabe et al., 2015; Stocker et al., 1997); moth: 60 glomeruli and 740 uPNs (Homberg et al., 1988); honeybee: 165 glomeruli and 800 uPNs (Brandt et al., 2005; Galizia et al., 1999; Rybak et al., 2010)), in the American cockroach each glomerulus is innervated by a single uPN (Ernst and Boeckh, 1983). Such an architectural difference should have an impact on the AL network coding properties, and might indicate species-specific adaptations to the environment, and distinct evolutionary strategies to efficiently encode olfactory inputs.

Recently, Nishino *et al*. showed an antennotopic distribution of pheromone-sensitive ORNs and their uPNs, which allows a spatial encoding of pheromone plumes along a single antenna (Nishino et al., 2018). Such an antennotopic organization could provide the means for encoding spatial information of complex olfactory environments. This becomes particularly relevant in dense dark shelters, where insects are required to rely heavily on smell. In such conditions, the ability to assess the spatial structure of a stimulus could provide pivotal information for odour-source localization. In addition, spatial separation can also aid in differentiating between homogeneous and heterogeneous mixtures, because in the latter, plumes have pockets of different odour quality, and thus can be used to identify different odorant sources (Nowotny et al., 2013; Stierle et al., 2013).

Here, by selectively labelling antennal lobe uPNs, we study how olfactory stimuli of different nature and concentration are encoded within the AL of a hemimetabolous insect. We selected similar and dissimilar odorants to probe how structural relatedness is encoded in the glomerular space. Furthermore, we extend the investigation of spatial coding to include also non-pheromone odour groups. We applied spatially confined stimuli to assess stimulus sensitivity along the antenna, and investigate how responses of AL output neurons (*i*.*e*. uPNs) change according to the stimulus nature, concentration, and location.

## Methods

### Animal preparation

Adult males of the American cockroach *Periplaneta americana (Linnaeus, 1758)* were kept on a 12/12 h light/dark cycle at 24°C and 65% humidity in the animal facility of the University of Konstanz (Germany). Male cockroaches were collected and briefly anesthetized with CO_2_ to facilitate handling. For **neuroanatomy**, a glass capillary coated with the fluorescent tracer Alexa Fluor 568 conjugated with 10 kDa dextran (all Alexa Fluor dyes were from Thermo Fisher Scientific, Waltham, MA, USA) was injected into the AL. For uPNs/mPNs double labelling, 10 kDa dextran-conjugated Alexa Fluor 488 and Alexa Fluor 647 were injected medio-ventrally to the medial calix of the mushroom body and medio-ventrally to the lateral protocerebrum, respectively. Dye-injected specimens were incubated in a humid chamber at 4°C for 24h. On the following day, brains were dissected, fixed in 4% paraformaldehyde (PFA) in phosphate buffered saline (PBS) for 3h at 4°C, dehydrated in ascending concentrations of ethanol, cleared in xylene, and mounted in DPX mounting medium (Sigma-Aldrich, St. Louis, MO, USA). Animal preparation for projection neurons’ **calcium imaging** was adapted from a protocol previously established in honeybees (Paoli et al., 2017; Sachse and Galizia, 2002). On the first day, male cockroaches were anesthetized and placed in a custom-build holder. The head was restrained with soft dental wax and a small window was opened in the head cuticle to expose the injection site. For uPN labelling, the tip of a glass capillary coated with 10 kDa dextran-conjugated Fura-2 (Thermo Fisher Scientific, Waltham, MA, USA) was injected ventro-medially to the medial mushroom body calix, where the ALT-I/II (Malun et al., 1993) travel along the mushroom body. For mPN labelling, the injection was directed to the ventro-medial side of the LP, where the outer-ALT formed by mPN axons enters the LP (Malun et al., 1993). All animals used in this study were labelled on the right side of the brain only. After dye injection, the head capsule was closed to prevent brain desiccation. On the following day, the antennal lobe was exposed to allow optical access, and the brain was covered in transparent two-component silicon (Kwik-Sil, WPI, Sarasota, FL, USA). In this procedure, the degree of labelling depends on the precision of the injection site and on the amount of dye loaded on the glass capillary. Hence, a variable level of PN labelling may result. To investigate the topology of OSN axon terminals, selective degeneration of OSNs was induced by amputating distal antennal segments. The flagellum of a briefly anesthetized cockroach was left intact or cut at different proximal-distal sections (25%, 50%, 75%, or 100% amputation). Cockroaches were then kept in a dark cage with food and water supplied *ad libitum* for a seven-day recovery period. During this time, all OSNs originating in the amputated part of the antenna would degenerate, and their axon terminals disappear (Distler and Boeckh, 1997; Nishino et al., 2018). After a week, the antennal nerve was cut at the scape and placed onto a glass capillary loaded with an aqueous solution of 10 kDa dextran-conjugated Alexa Fluor 555. In parallel, uPNs were labelled as described above with 10 kDa dextran-conjugated Alexa Fluor 488. Projection neurons labelling was used as reference to quantify glomerular volume even after partial degeneration of the OSN terminals. Dye-injected specimens were incubated in a humid chamber at 4° C for 24 h, after which the brain was dissected, and processed for confocal microscopic observation. Thirty-seven specimens were prepared for this analysis.

### Neuroanatomy

Whole brain sections (**Fig. 1A, Movie 1**) were acquired with a LSM 5 PASCAL laser scanning confocal microscope (Carl Zeiss, Jena, Germany) at a resolution of 1.16 µm/pixel (x, y) and 2 µm z-intervals using a 10x dry objective (Zeiss Plan-Apochromat 10x/0.45). Antennal lobe optical sections with uPNs/mPNs labelling (**Fig. 1C**) were acquired with an LSM 510 laser scanning confocal microscope (Carl Zeiss, Jena, Germany) at a resolution of 0.77 µm/pixel (x, y) and 3 µm z-intervals using a 10x water-immersion objective (Zeiss C-Apochromat 10x/0.45w); antennal lobe optical sections with uPNs labelling (**Fig. 1D**) were acquired at a resolution of 0.54 µm/pixel (x, y) and 1 µm z-intervals using a 20x water-immersion objective (Zeiss W-PlanApochromat 20x/1.0); antennal lobe optical sections for glomerular volume reconstruction (**Fig. 2**) were acquired at a resolution of 0.38 µm/pixel (x, y) and 3 µm z-intervals using a 20x water-immersion objective (Zeiss W-PlanApochromat 20x/1.0). ImageJ was used for image processing.

**Figure 1.**
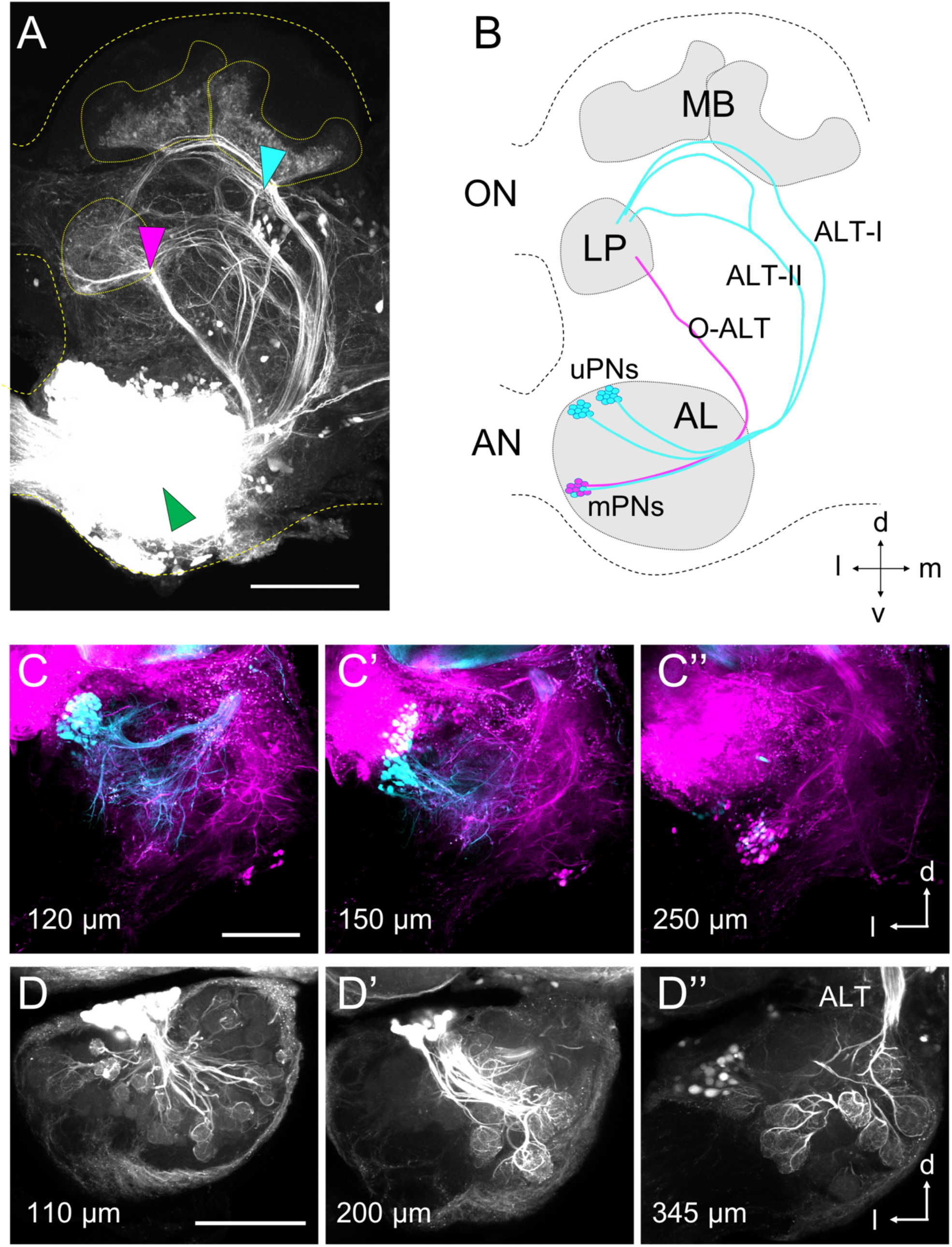
Antennal lobe projection neurons labelling. (**A**) Antennal lobe injection (green arrowhead) reveals the main PN tracts. Local injection ventro-medial to the medial MB calix (cyan arrowhead) labels uPNs; injection ventral to the LP (magenta arrowhead) labels mPNs. (**B**) Drawing of the main PN tracts. AL, antennal lobe; MB, mushroom body calyces; ALT, antennal-lobe tract; LP, lateral protocerebrum; ON, optic lobe nerve; AN, antennal nerve. (**C**-**C“**) Double labelling for uPNs (cyan) and mPNs (magenta), at different focal depths. (**D**-**D“**) Uniglomerular PN labelling at different focal depths. Numbers indicate the acquisition depth along the AL antero-posterior axis. Scale bars, 200 µm.

**Figure 2.**
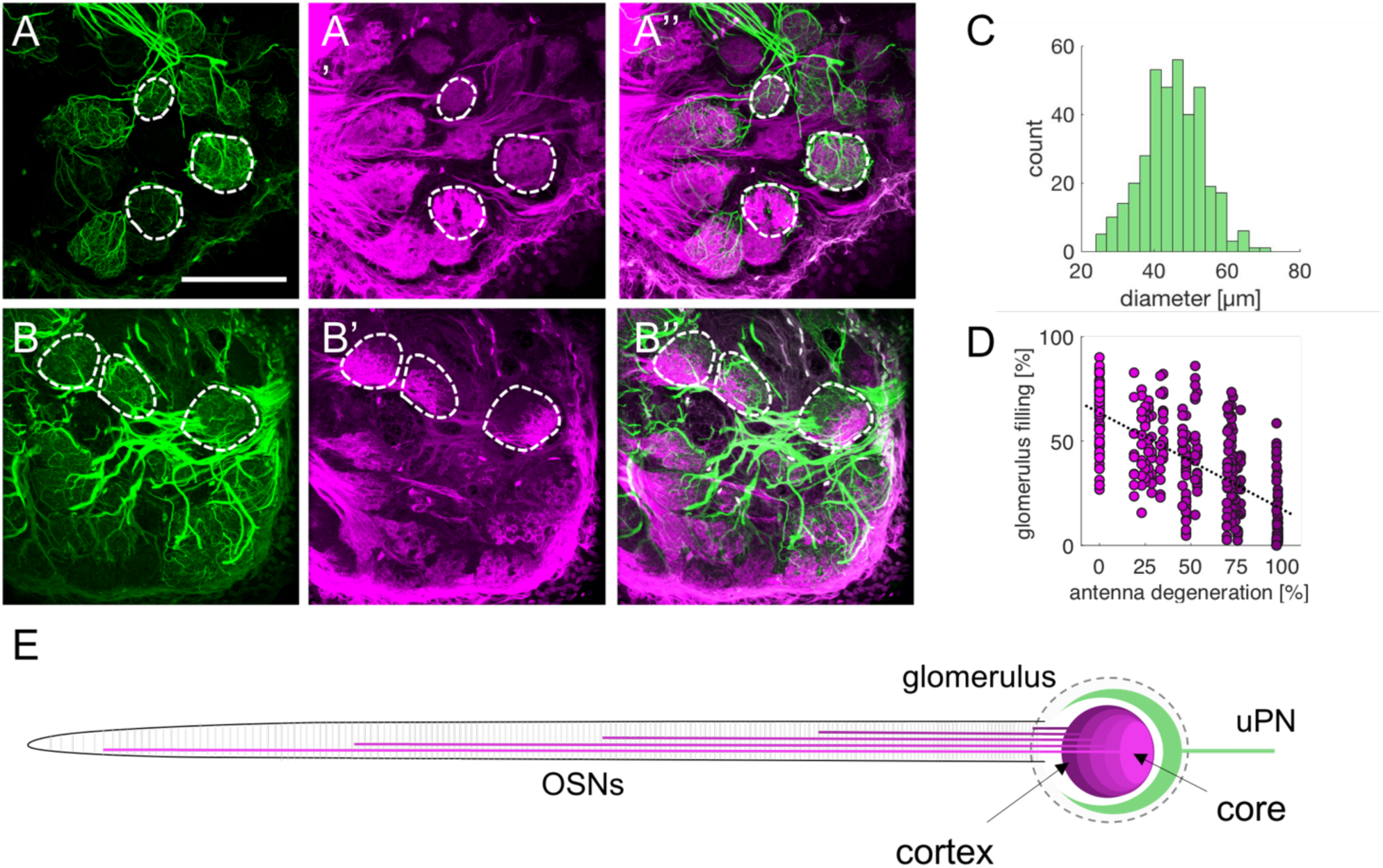
Selective degeneration of olfactory sensory neurons revealed an antennotopic innervation pattern. (**A**,**B**) Optical section through an AL labelled for OSNs (magenta) and for uPNs (green). In (**A**), the antenna was left intact, whereas in (**B**) 50% of the antenna was amputated before OSNs tracing. Three representative glomeruli are circled by a dashed line. Scale bar, 200 µm. (**C**) Quantification of glomerular diameter (mean ± s.d. = 45 ± 8 µm, *n* = 369 glomeruli from 37 antennal lobes) (**D**) Quantification of OSNs glomerulus filling upon selective antennal nerve degeneration. Increasing portion of degenerated antennal nerve induces progressive loss of innervation (linear regression analysis, R^2^ = 0.43). (**E**) Schematics of glomerular innervation topology based on OSNs origin along the antenna according to our results and previous reports (Nishino and Mizunami, 2007; Nishino et al., 2018).

### Calcium imaging

Calcium imaging analysis of AL projection neurons was performed at a wide field fluorescence microscope (BX51WI, Olympus, Tokyo, Japan) equipped with a 10x water immersion objective (Olympus UM Plan FI 10x/0.30w). Images were acquired with a SensiCam CCD camera (PCO AG, Kelheim, Germany) with a 4×4 binning configuration resulting in 120×160 pixel sized images (corresponding to 450×600 µm, with a pixel size of 3.75×3.75 µm). Recordings were performed at 5 Hz (for odour response maps, **Fig. 4**) or at 25 Hz (for measuring responses to localized stimuli, **Fig. 5**) using a TILLvisION acquisition system (TILL Photonics, Graefelfing, Germany). A LED system equipped with a 340 and a 385 nm LED (Omicron-Laserage Laserprodukte GmbH, Rodgau-Dudenhofen, Germany) was employed as light source, which was directed onto the cockroach brain via a 410 short-pass filter and a 410 dichroic mirror. Emitted light was filtered through a 440 long-pass filter. Videos were exported and processed in Matlab (MathWorks Inc, Natick, MA, USA). Calcium imaging of the macroglomerulus was performed at a LSM 510 laser scanning microscope equipped with a Ti:Sapphire two-photon laser (Chameleon Ultra, Coherent, CA, USA). Calcium sensitive dye Fura-2 was excited with 750 nm pulses and emitted light was filtered through a 650 nm short-pass filter. The area corresponding to a central section of the macroglomerulus was scanned at a 5 Hz frequency and with 2 µm/pixel (x, y) spatial resolution.

**Figure 3.**
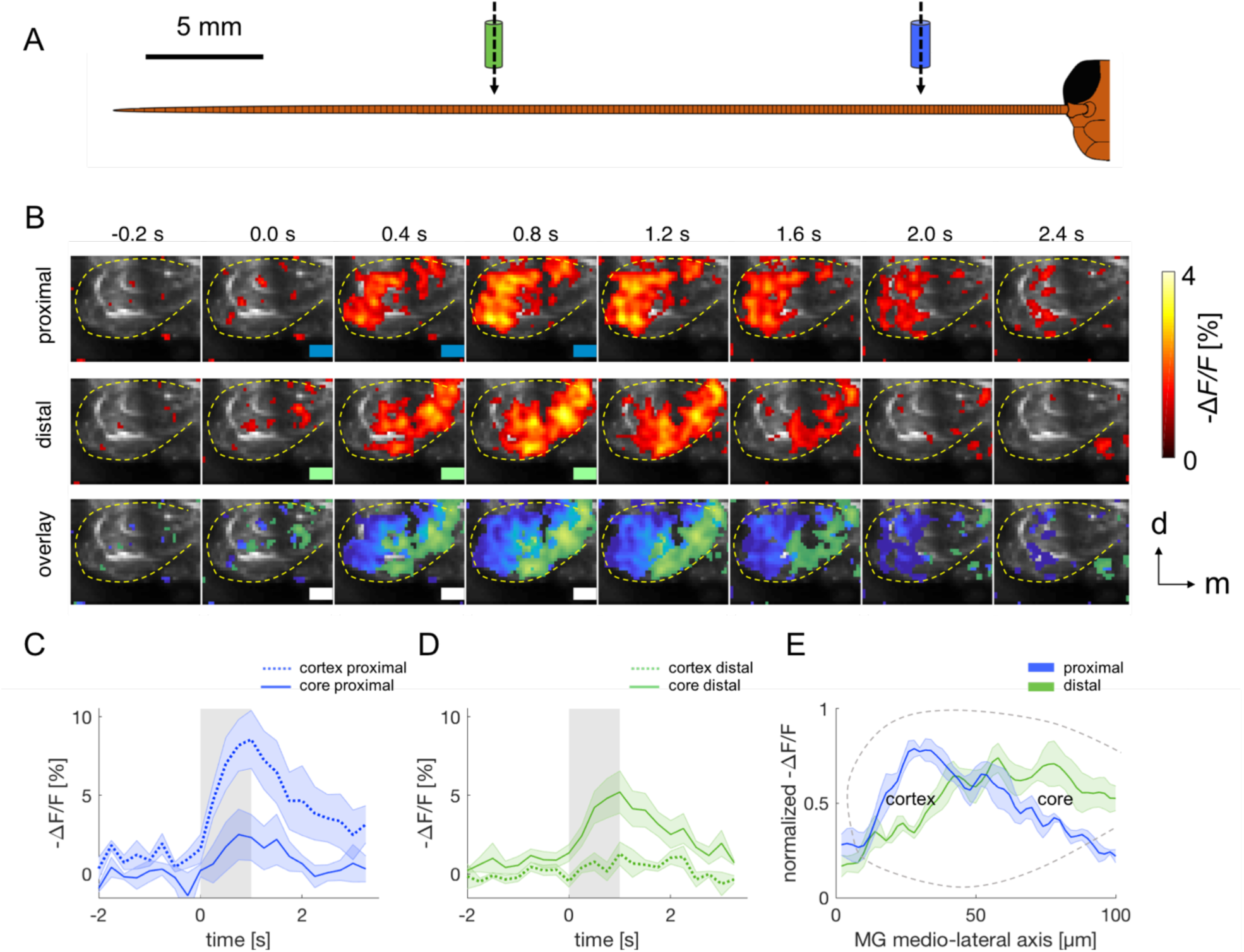
Antennotopic odour response in the macroglomerulus. (**A**) Schematic representation of localized stimulation experiment. Proximal stimulation (blue, 6 mm from scapus), distal stimulation (green, 24 mm). (**B**) Time sequence of a representative MG response to a proximal and a distal periplanone-B stimulation. The yellow dashed line delimits MG; green/blue rectangles indicate distal/proximal stimulus delivery; scale bar, 50 µm. (**C-D**) Mean response (± s.e.m.) of MG core (continuous line) and cortex (dotted line) to a proximal (**C**) and a distal (**D**) stimulation. (**E**) Mean evoked responses (± s.e.m.) along the longitudinal MG axis for proximal and distal stimulation, where position 0 µm corresponds to the most lateral side and position 100 µm to the most medial side of the MG. See **Fig. S1** for the original traces.

**Figure 4.**
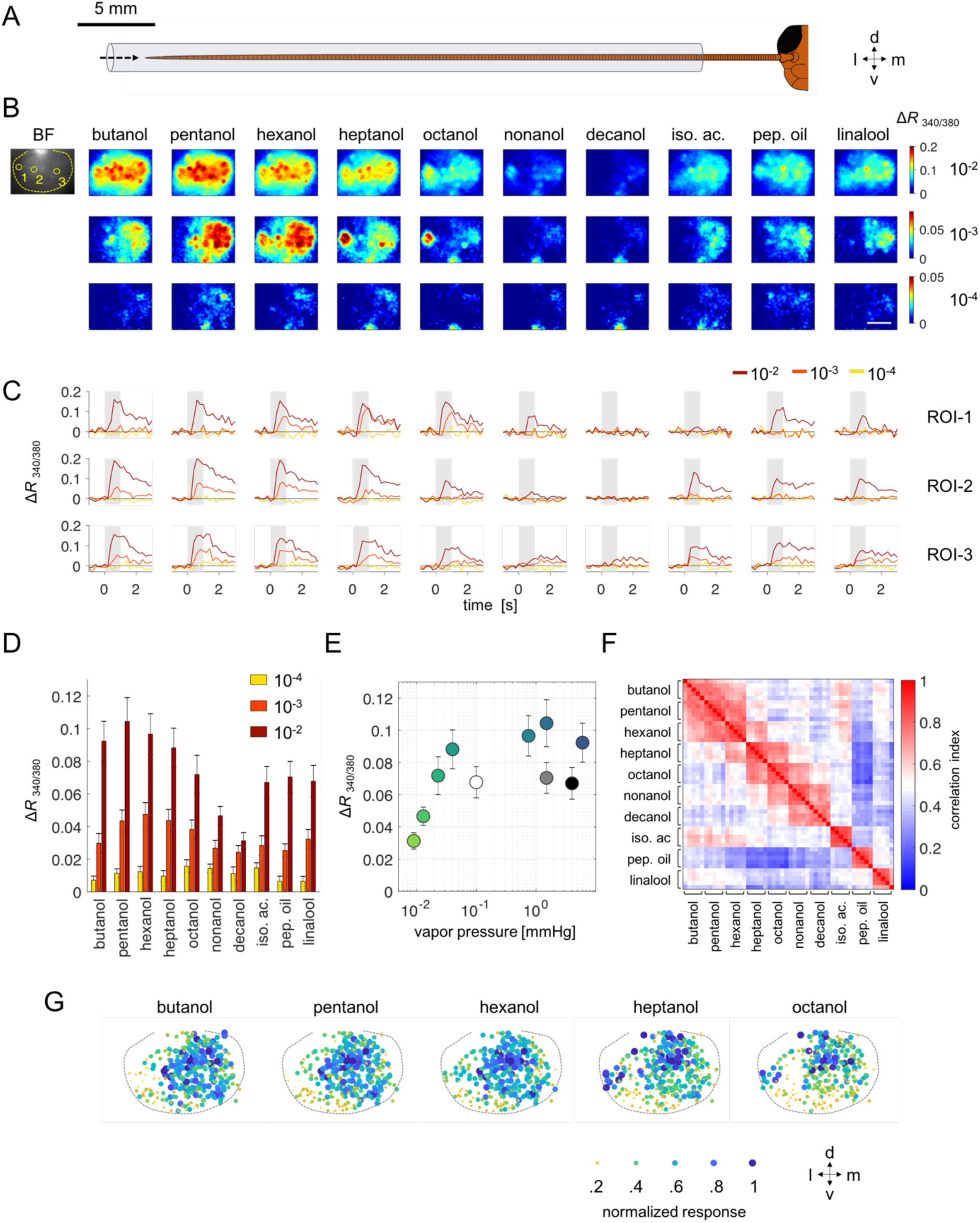
Common odorants evoke stimulus-specific combinatorial patterns in the antennal lobe. (**A**) Schematic representation of localized stimulation experiments. (**B**) Representative panel of evoked responses in one individual for 10 odorants at 3 dilutions. A dashed line in the bright field (BF) image indicate the AL. Scale bar, 200 µm. (**C**) Temporal response profiles of the three regions of interest (ROI-1,-2,-3) indicated with yellow circles in the bright field image of panel (**B**). Colours refer to different dilutions; grey bars in the plots indicate stimulation window. (**D**) Mean ± s.e.m. of maximum responsive region for all odorants and dilutions (*n* = 11 cockroaches). (**E**) Mean ± s.e.m. of maximum responsive regions for the 10^−2^ dilution are plotted versus odorant vapour pressure. (**F**) Correlation analysis among glomerular response vectors of five repetitions of 10 odorants at 10^−3^ dilution. Odorants were delivered in a pseudorandomized order. Mean correlation map for 9 animals. (**G**) All odour-induced responses from 9 cockroaches were normalized within each animal and plotted onto a reference antennal lobe. Normalized intensity of the response is encoded by marker’s color and size.

**Figure 5.**
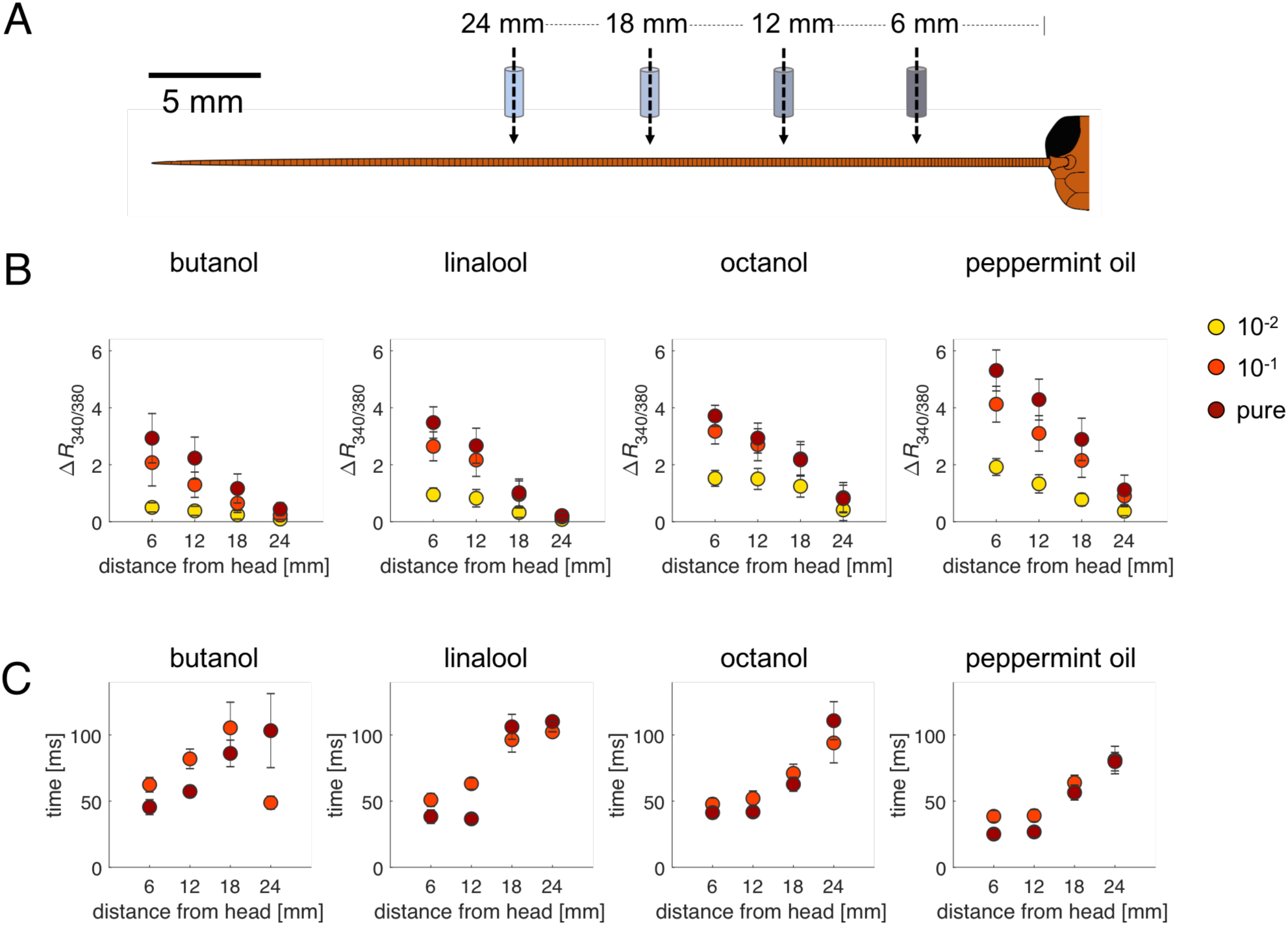
Analysis of response latency and intensity of spatially localized olfactory stimuli. (**A**) Schematic representation of localized stimulation experiments. (**B**) Mean response intensity ± s.e.m. relative to the strongest responsive regions of the AL for four stimulus locations and three dilutions. (**C**) Mean latency ± s.e.m. of the same strongest responsive regions for four stimulus locations and two dilutions. *n* = 14 cockroaches.

### Odorant preparation

Odorants used were 1-butanol (CAS 71-36-3, cat.num. 19422-5ML), 1-pentanol (CAS 71-41-0, cat.num. 77597-1ML-F), 1-hexanol (CAS 111-27-3, cat.num. 73117-1ML-F), 1-heptanol (CAS 111-70-6, cat.num. 72954-1ML-F), 1-octanol (CAS 111-87-5, cat.num. 95446-5ML-F), 1-nonanol (CAS 143-08-8, cat.num. W278904-SAMPLE-K), 1-decanol (CAS 112-30-1, cat.num W236500-SAMPLE-K), linalool (rac., CAS 78-70-6, cat.num. L2602-100G), isoamyl acetate (CAS 193-12-2, cat.num. 79857-5ML), peppermint oil (CAS 8006-90-4, cat.num. 77411 - 25 ml). All odorants were purchased at Sigma-Aldrich (St. Louis, MO, USA) in the highest purity available. Odorants were used as pure or diluted in mineral oil (CAS 8042-47-5, cat.num. 124020010, Acros Organics, Thermo Fisher Scientific, Waltham, MA, USA). Dilutions were prepared in 5 ml (for experiments with the PAL autosampler), or in 1 ml mineral oil (for experiments of spatially localized stimulation performed with the fast olfactometer) in 20 ml glass vials, covered with nitrogen (Sauerstoffwerk Friedrichshafen GmbH, Friedrichshafen, Germany) to prevent oxidation, and sealed with Teflon septum (Axel Semrau GmbH, Sprockhövel, Germany). Synthetic periplanone-B (Kuwahara and Mori, 1990) was distributed by Prof. Mark Willis to H.N. (Case Western Reserve University) diluted in hexane (CAS 110-54-3).

### Odorant delivery

An automatic multi-sampler for gas chromatography (Combi PAL, CTC Analytics AG, Zwingen, Switzerland) was adapted for olfactory stimulation. Two 1-ml odour pulses were injected at 4 and 12 s with an injection speed of 1 ml/s into a continuous flow of 1 ml/s purified air. The stimulus was directed towards the antenna via a Teflon tube (inner diameter, 0.87 mm; length, 40 cm), and the antenna was inserted into the terminal portion of the tube to ensure similar stimulation conditions across animals. The first stimulus reached the antenna with ∼800 ms delay due to delays in the sampler and to the air flow. Hence, first stimulus onset was determined as 4.8 s. An inter-trial interval of two minutes was kept between odorants, during which the syringe was flushed with clean air. For each animal, response to mineral oil was tested as negative control. At the end of a stimulation cycle, odour response to the first odorant was tested once again to assess the stability of odour responsiveness. After each set of odorants, the syringe was washed with *n*-pentane (Merk KgaA, Darmstadt, Germany), heated to 44°C, and flushed with clean air.

For the spatially localized stimulations, we employed a fast odour-delivery device with rise time <3 ms (Raiser et al., 2017) controlled by a stimulus controller (cRIO-9074 combined with IO module NI-9403, National Instruments, Austin, TX, USA) coupled to a LabVIEW software (National Instruments) custom-written by Stefanie Neupert. The outlet of the olfactometer, a glass tube with a 2 mm inner diameter, was placed at a right angle to the antenna at <1 mm from the antenna at different proximal-to-distal positions (*i*.*e*. 6, 12, 18, 24 mm from the scapus). Given a total antennal length of 40-to-50 mm, such experimental configuration would stimulate ∼5% of the antennal length. For each odorant and concentration, three stimuli of 100 ms with 10 s inter-stimulus interval were applied. Odorants were given in a pseudorandomized order; concentrations were created with dilution between 10^−2^ and undiluted (pure) odorants. The air carrier flow was set to 1 l/min and the valve-controlled flow was set to 300 ml/min, for a total constant flow of 1.6 l/min (Raiser et al., 2017). Odour arrival, calibrated with a photoionization detector, was 120 ms after trigger onset, corresponding to a 3-frame delay at 25 Hz acquisition rate.

For periplanone-B stimulation, we adopted a simplified version of the fast-odour delivery device described above to reduce tubing length. A filter paper loaded with 10 µl of purified periplanone-B was loaded immediately upstream of a three-way solenoid valve (Lee Products Ltd, Gerrards Cross, UK) controlled by the previously described LabVIEW software. In non-stimulating conditions, the valve received an input of clean air, directed into an air-carrier tube (length 40 mm, ID 3.2 mm, Tygon-tubes 2001 ultra-pure, Carl Roth, Karlsruhe, Germany), whose outlet was placed ∼1 mm from the antenna and perpendicular to it either in a proximal (6 mm) or distal (24 mm) position (**Fig. 3A**). The air carrier flow was set to 800 ml/min and the valve-controlled flow was set to 150 ml/min, for a total constant flow of 950 ml/min. During stimulation the valve switched from clean air to stimulus. For each experiment, we delivered three stimuli of 1 s at *t* = 2, 10, and 18 s.

### Data analysis

Glomerular 3D reconstruction and volumetric analysis (**Fig. 2**) were conducted with Amira Software (Thermo Fisher Scientific, Waltham, MA, USA). Analysis of glomerular filling of olfactory sensory neurons was conducted in blind: a non-informative code was assigned to each brain sample, and both confocal imaging and 3D reconstruction were conducted without knowing the extent of antenna amputation. Regions of maximal response in the MG (**Fig. 3**) were automatically defined based on the global response maxima to proximal and distal stimulations. In brief, average response maps to proximal and distal stimuli were segmented using a multilevel threshold (the Otsu algorithm, implemented in Matlab, MathWorks Inc, Natick, MA, USA) to detect the highest activity region >50 pixels (∼100 µm^2^). Since in all tested preparations these were located in the outer and inner part of the MG respectively, we labelled them as “cortex” and “core”. At the wide-field fluorescence microscope, we used a ratiometric protocol for calcium imaging (Fura-2 has excitation wavelengths of 340 and 380 nm and emission of 510 nm). Ratios of 340 to 380 nm signals were calculated (*R*_340/380_), and offset was removed by subtracting the mean signal before odour stimulation (Δ*R*_340/380_). To construct odour response maps, we considered the mean activity between 0.6 to 1 s after stimulus onset. Response to the solvent (mineral oil) was subtracted from all stimulus-induced signals. For quantifying response maps across the entire AL (**Fig. 4**), 42 ± 2 (mean ± s.e.m., range 29 to 50, *n* = 9 cockroaches) responsive areas of 5-by-5 pixels (corresponding to 19-by-19 µm) were hand-selected to construct stimulus specific response vectors. Correlation analysis across response vectors of same and different stimuli were performed to assess stimulus specificity of the elicited response and odour similarity (**Fig. 4F**). To assess spatial representation of alcohols with increasing chain length, responses from all animals (*n* = 9) were pooled together onto the same meta-antennal lobe. To focus on relative intensity and spatial distribution, all responses from each animal/odorant combination were normalized as *resp*_*g_norm,a,o*_ = (*resp*_*g,a,o*_ – *resp*_*min,a,o*_)/(*resp*_*max,a,o*_ – *resp*_*min,a,o*_), where *resp*_*g,a,o*_ is the response intensity in a specific area, *resp*_*min,a,o*_ and *resp*_*max,a,o*_ are the minimum and maximum response intensity in the antennal lobe of animal *a* for odorant *o*. Normalized responses to alcohols were plotted into a standard antennal lobe according to their original coordinates and size in a colour coded map. For the analysis of responses to stimuli at different locations along the antennae (**Fig. 5**), amplitudes were calculated as mean response intensity between 120 and 280 ms after stimulus onset. To determine response latency, response curves were fitted with the sigmoidal function f(*t*) = *a* / (1+e^- *b*(*t*-*c*)^) +*d*, where *a* is the amplitude of the fitted curve, *a*×*b*/4 is the slope of the sigmoid, *c* is the time of 50% response, and *d* is the minimum value of the sigmoid. Response latency was defined as the time point along the fitted curve where intensity value crossed a threshold of 2σ of the pre-stimulus activity. Calcium imaging data analysis was conducted with custom made Matlab scripts (MathWorks Inc, Natick, MA, USA).

### Statistical analysis

Relationship between antennal degeneration and OSNs glomerular filling (**Fig. 2D**) was assessed with a linear regression analysis. Responses to proximal/distal stimuli in the MG core/cortex (**Fig. 3E**) were analysed with paired Student’s *t*-test. The effect of odorant, dilution, and stimulus location on signal intensity and latency (**Fig. 5**) was assessed with a 3-way analysis of variance. Travelling speed of neural activity along the antenna was calculated for each animal (*n* = 14) by fitting all response latencies for the different odorants, dilutions and locations with a linear regression.

## Results

### Uniglomerular projection neurons in the AL: morphology

Antennal lobe injection with the fluorescent tracer Alexa Fluor 546 revealed the multiple AL projection neuron tracts (**Fig. 1A-B**, green arrowhead; **Movie 1**). The neuroanatomy of ALTs is coherent with previous observations (Malun et al., 1993), and was used to guide us in performing selective retrograde labelling, in a protocol adapted from other insect species, *e*.*g*. honeybees (Sachse and Galizia, 2002), or ants (Zube et al., 2008). A localized tracer injection into the medial tracts ALT-I/II medio-ventrally to the mushroom body calices (**Fig. 1A-B**, blue arrowhead) allowed for uPNs population backfilling. Their somata group into two main clusters, type I (**Fig. 1C**) and type II (**Fig. 1C’**), located on the antero-dorsal side of the AL (Strausfeld and Li, 1999; Watanabe et al., 2017). Alternatively, dye injection on the ventral side of the lateral protocerebrum (magenta arrowhead) allowed labelling of multiglomerular projection neurons (via the outer-ALT), whose somata localize postero-laterally with respect to the uPN clusters (**Fig. 1C”**). Here, we focused on uPNs labelling to investigate the basis of olfactory coding in a hemimetabolous insect. After tracer injection, the cell body clusters and their dendritic arborizations within the glomeruli could be easily identified (**Fig. 1D, Movie 2**). Observation of uPNs labelling confirmed previous studies indicating that each glomerulus is innervated by only one uPN, with the exception of the macroglomerulus (Schaller, 1978; Watanabe et al., 2017).

Given that previous works had focussed on the macroglomerulus only, we investigated antennotopic innervation in ordinary glomeruli. By removing a distal portion of the antenna (at 0, 25, 50, 75, or 100% of its length), we selectively induced neurodegeneration of the ORNs originating in the missing part. After seven days, we bulk labelled all remaining sensory axons and evaluated the decrease in OSN-innervated area within randomly selected glomeruli using 3D reconstruction of confocal imaging (**Figure 2**). Average intact glomerulus diameter was 45 ± 8 µm, measured from reconstruction of PNs staining in *n* = 369 glomeruli (**Fig. 2C**). From that, using uPNs labelling as counterstaining to visualize the intact glomerulus, we could assess OSNs innervation volume after partial OSN terminals degeneration. We found that the glomerular volume occupied by OSN terminals was reduced proportionally to the percentage of missing antenna (**Fig. 2D**) and that OSNs originating in different antennal segments innervated different glomerular compartments (linear regression analysis, R^2^ = 0.43). Interestingly, despite the difference between the number of OSNs in ordinary glomeruli and in the MG, their innervation pattern was conserved (Nishino et al., 2018): OSNs originating on the distal portion of the antenna innervated the portion more proximal to the projection neuron’s innervation site, whereas OSNs hosted in proximal antennal segments innervated the glomerular region further from the uPNs entry site. Despite of this consistent pattern, the detailed distribution showed a high variability across glomeruli **(Fig. 2D)**, indicating that the distribution of receptor types along the antennal flagellum might differ for different receptor types.

### Antennotopic odour response in the macroglomerulus

Antennotopic glomerular innervation provides the means to relay spatial information from along the antennal flagellum to the central nervous system. The male macroglomerulus responds to the main component of the female sex pheromone periplanone-B, and is innervated by multiple spatially-segregated uPNs (Hösl, 1990; Nishino et al., 2018). We used this arrangement to test whether the spatial innervation of OSNs leads to spatial information in uPNs, and how odour locations are mapped by the MG uPNs. We used confocal imaging of Fura-2 loaded projection neurons and compared proximal and distal stimulations (6 and 24 mm from the head, **Fig. 3A**). We found that, consistent with the anatomical observations, distal stimulations elicited calcium responses in a more medial portion of the MG (“core”), while proximal ones activated more lateral (“cortical”) regions (**Fig 3B-E**) (paired Student’s *t*-test, *p* < 0.05, *n* = 4). In addition, proximal stimuli induced stronger responses, whereas distal stimuli consistently elicited weaker ones (**Fig. 3C-E**; see **Fig. S1** for individual traces).

### Uniglomerular projection neurons’ calcium imaging

Next, we investigated how common odorants are represented in the cockroach’s antennal lobe. Medial PN tracts were retrogradely filled with the calcium-sensitive dye Fura-2. Successful labelling resulted in bright fluorescence of the cell bodies and dendritic arborizations (**Fig. 1D, Movie 2**), and allowed calcium imaging during olfactory stimulations for several hours. Before stimulations, spontaneous changes in calcium amplitudes could be observed in discrete areas, comparable in size and shape to individual glomeruli. During stimulation (schematics in **Fig. 4A**), fluorescent changes were observed in these discrete areas, as well as in larger portions of the AL, suggesting that in some instances, clusters of neighbouring glomeruli were activated simultaneously (**Movie 3**). Because it was not possible to reliably identify individual glomeruli across animals, we compared odorant and concentration-induced response maps within individuals, and extracted general coding features from an across-individual comparison. Eleven cockroaches were tested with a panel of ten odorants at three dilutions (10^−2^, 10^−3^, and 10^−4^). Response intensity was stimulus-dependent, with long-chain alcohols (1-nonanol and 1-decanol) inducing much weaker responses than shorter-chain ones (1-butanol to 1-heptanol). Other tested compounds (1-octanol, linalool, isoamyl acetate and peppermint oil) induced concentration-dependent robust responses, with smaller amplitudes than for short-chain alcohols. **Fig. 4B** provides a representative panel of odour response maps. Each image (600-by-450 µm) shows the anterior view of the right antennal lobe. Within the responsive area, spherical spots of activity of ∼45 µm in diameter were clearly visible, likely corresponding to activated AL glomeruli. Different concentrations were tested to investigate the sensitivity of the cockroach’s olfactory system, showing clear stimulus-induced responses for high and medium dilutions, whereas no response beyond the negative controls were observed for 10^−4^ dilutions (**Fig. 4B, C**). This effect was consistent across all tested odorants. For all odorants, spatial response patterns at lower dilutions were confined to smaller areas. Accordingly, different glomeruli had odorant-specific concentration response curves. For instance, as shown in **Fig. 4C**, ROI-1 and ROI-3 responded with similar intensity to butanol at 10^−2^, but while ROI-3 responded with 10% to 1-butanol 10^−3^ stimulation, ROI-1 was silent at this concentration. Similarly, linalool 10^−2^ triggered a comparable response both in ROI-2 and ROI-3, whereas at 10^−3^ its induced activity in ROI-3 showed half of the response to the highest concentration, while ROI-2 was already non-responsive.

For each odour response map, we considered the area with the strongest activity to estimate odorant and concentration sensitivity across animals. For each odorant/dilution pair, we computed a mean response value (*n* = 11 animals). Overall, cockroaches’ uPNs displayed a response of 0.07-0.11 Δ*R*_340/380_ in response to odour stimulations at 10^−2^ (a lower response for long-chain alcohols), 0.02-0.05 to intermediate dilution, and 0.01-0.02 to 10^−4^ dilutions (**Fig. 4D**). The number of molecules in a stimulus scales, to a large degree, with its vapor pressure. **Fig. 4E** illustrates the relationship between odorants vapor pressure and AL activity. We observed a decrease in responsiveness for long-chain alcohols, which have the lowest vapour pressure of the tested odorants. However, for the other odorants we found comparable response intensities to molecules with very different vapor pressures. For instance, response elicited by 1-octanol and 1-butanol, whose vapor pressures differ by 260 fold (6 *vs* 0.023 mmHg), were comparable. Similarly, isoamyl acetate, peppermint oil and linalool, despite of their different vapour pressures (0.1, 1.5 and 4.0 mmHg, respectively) evoked comparable response intensities (**Fig. 4E**). Thus, at least for some stimuli, the olfactory system appears to compensate for vapor pressure, *i*.*e*. for expected molecule density in the odour plume.

Physiological odour similarity can be assessed by measuring the correlation coefficient between odour response maps (Guerrieri et al., 2005). With this purpose, we exposed a new group of cockroaches (*n* = 9) to five repetitions of ten odorants at an intermediated dilution (10^−3^) in a pseudorandomized order. For each animal, we constructed stimulus response vectors considering all functional units activated by any of the odorants. We obtained fifty vectors from five repetitions of ten odorants, and calculated a correlation matrix across all vectors to quantify odour response similarity. We found that repetitions of the same stimulus were highly correlated (**Fig. 4F)**, whereas different odorants showed different degrees of similarity/dissimilarity. Interestingly, chemically similar alcohols showed higher correlation values than chemically dissimilar ones (based on carbon chain length). How is the “chain length” parameter represented across the cockroach’s glomerular space? To answer this question, we mapped the responsive areas from all individuals and overlaid them onto a common reference antennal lobe. Then, we analysed the spatial distribution of the observed responsive glomeruli for the chosen alcohol series, with the exclusion of 1-nonanol and 1-decanol because of their generally low responsiveness (**Fig. 4G**). Notably, a cluster of strongly responsive glomeruli in the central part of the antennal lobe was present for all alcohols. However, short-chain alcohols tended to activate more central/medial regions of the AL, whereas with increasing chain length, the responsive area shifted from centro-medial to centro-lateral.

### Representation of spatially confined stimuli

With the previous set of experiments, we considered the antenna as a whole, and stimulated the entire array of olfactory sensory neurons. However, OSNs that express the same olfactory receptor maintain an antennotopic organisation within the glomeruli (**Fig. 2**). To investigate how spatially confined stimuli are encoded in the AL, we used short stimulations and a narrow olfactometer outlet placed perpendicularly to the antenna at different locations, *i*.*e*. at 6, 12, 18, 24 mm from the scapus (olfactometer inner diameter, 2 mm; distance from antennal surface, < 1 mm, see schematics in **Fig. 5A**). We selected 24 mm as the most distal position because more distal stimuli rarely produced detectable responses. For this experiment, we employed four odorants, eliciting strong and clear responses: a short- (1-butanol) and a long-chain alcohol (1-octanol), linalool, and peppermint oil. In addition, 3 dilutions were used: pure odorants, 10^−1^, and 10^−2^. This experimental design required lower odorant dilutions than before because of its high spatial and temporal confinements. For each odour/dilution/position/animal combination we considered the region of the AL with the strongest stimulus-induced response, and analysed its response intensity and latency. Response intensity (**Fig. 5B**) depended on the odorant (3-way ANOVA, *df* = 3, *F* = 19.79, *p*<0.01), concentration (*df* = 2, *F* = 44.28, *p*<0.05), and stimulus location (*df* = 3, *F* = 48.75, *p*<0.01), with proximal stimuli generating stronger responses than distal ones. Notably, significant interactions were found between factors concentration and position (*df* = 6, *F* = 4.33, *p*<0.01), underlining the fact that a significant effect of concentration on the elicited response depends on the position effect (*i*.*e*. at distal locations, responses are weaker and concentration-dependent differences cannot be observed). To quantify response latency (**Fig. 5C**), we fitted the rising part of the response profile with a sigmoidal curve and considered as response onset the timepoint of crossing a threshold of two standard deviations from baseline activity. For this, we only used the two strongest concentrations, whose responses were robust enough to provide reliable data fitting, and observed that latency increased with distance from the head. In this case the sole factor influencing response latency was stimulus location (*df* = 3, *F* = 7.76, *p*<0.01), while odorant (*df* = 3, *F* = 2.04, *p* = 0.11) and concentration (*df* = 1, *F* = 0.51, *p* = 0.47) had no significant effect. We performed a linear regression for stimulus location to assess propagation speed along the antenna. Mean OSNs’ conduction velocity (± s.e.m.) was 0.37 ± 0.04 m/s, ranging from 0.19 to 0.58 m/s across animals (*n* = 14 preparations), with a delay of 30 ± 16 ms between the stimulation of the most proximal antennal section and the first detectable evoked activity in the AL projection neurons. Considering that a proximal stimulation elicits a OSNs activity at ∼7 mm from the AL, and that such activity travels along the antennal nerve at 0.37 m/s, it will reach the AL with a latency of ∼19 ms. Our observation that a proximal stimulation induces AL activity with ∼30 ms time delay suggests that the average lag between the OSNs input signal and a detectable calcium activity in the AL output neurons is approximately 11 ms. Using the same odour delivery device used in our study, Egea-Weiss *et al*. were able to quantify OSNs response latency in the fruit fly (Egea-Weiss et al., 2018), showing that OSNs response time may be as fast as 3 ms from odour arrival. In addition, it has been reported that also other insects (*i*.*e*. the hissing cockroach, the locust, the silk moth, and the honeybee) have a OSN response latency of a few ms after stimulous arrival (Szyszka et al., 2014). Assuming that the American cockroach OSNs have a similar response dynamics, our analysis suggests that an olfactory input needs approximately 8 ms to be processed within its AL network.

## Discussion

The olfactory circuit of *P. americana* has been extensively investigated thanks to its robustness and accessibility for long-term electrophysiology and neuronal tracing (Ernst and Boeckh, 1983; Hösl, 1990; Husch et al., 2009; Malun et al., 1993; Salecker and Boeckh, 1996). Hence, being amongst the best described olfactory systems, it provides an excellent model for understanding which olfactory coding rules apply to hemimetabolous insects. In addition, the cockroach’s extremely long antennae, both in absolute and relative terms, offer the opportunity to study if and how spatial information of odorants is encoded in the antennal lobe. With this in mind, we set here to assess how stimulus nature, concentration, and spatial location are mapped within the cockroach olfactory system.

### Coding similarity of holometabolous and hemimetabolous insects

In hemimetabolous insects, different solutions exist for the glomerular organization of the antennal lobe: the American cockroach has a comparatively small number of large glomeruli, and uniglomerular projection neurons (Watanabe et al., 2010), which is similar to the arrangement in most holometabolous insects studied so far (*e*.*g*. fruit flies, honeybees, moths); locusts, however, have a high number of microglomeruli, and only multiglomerular projection neurons (Hansson et al., 1996; Moreaux and Laurent, 2007). The question arises, therefore, whether the cockroach AL has a functional organization that is similar to the one of holometabolous insects, or rather to the more closely related locust. Selective labelling of AL projection neurons was initially developed for the honeybee (Sachse and Galizia, 2002), and later adapted to other species. For the first time, we were able to transfer this method to a hemimetabolous insect, thus opening the way for comparative investigations of odour coding in holo-*vs*. hemimetabolous insects. This will allow to define essential common features in the ways volatile molecules are detected and processed: differences in architecture and physiology of these two insect groups would indicate on alternative evolutionary mechanisms for probing an olfactory environment; alternatively, common mechanisms would indicate a likely common origin of the underlying architecture.

One of the advantages of uPNs backfilling is the possibility to monitor a major part of the AL population at once. This allows the construction of broad response maps to study how different olfactory stimuli are spatially represented in the antennal lobe. A further advantage of this method in the American cockroach is that in this species each glomerulus is innervated by a single uPN (Ernst and Boeckh, 1983). As a consequence, all measured glomerular responses reflect the activity of exactly one neuron. Thus, differences in signal strength or quality across glomeruli cannot be attributed to the activation (or labelling) of a different number or proportion of neurons. Successful staining resulted in the clear labelling of the uniglomerular PNs, with their somata clusters in the dorso-lateral region of the antennal lobe (**Fig. 1, Movie 2**).

Calcium imaging analysis allowed us to measure how olfactory stimuli are represented in the AL within and across individuals. In agreement with the logic of a combinatorial code - described in holometabolous insects such as moths (Carlsson et al., 2002; Galizia et al., 2000), honeybees (Sachse et al., 1999), and fruit flies (Fiala et al., 2002; Silbering et al., 2008) - also in *Periplaneta*, each odorant induces the selective activation of a subset of the AL glomeruli. Within a single individual, glomerular response maps are stimulus-specific and reproducible, as shown by our correlation analysis (**Fig. 4F**). Testing different odorant dilutions revealed a concentration dependency of the elicited activity both in terms of intensity (stronger stimuli induced stronger responses), and responsive areas (stronger stimuli activated more glomeruli). In our analysis, we did not observe areas of the antennal lobe responsive to low but not to high odorant concentrations. Studies on the honeybee (Sachse et al., 1999) and on the fruit fly (Couto et al., 2005) have shown that chemical properties such as the alcohol chain length or functional groups can be encoded by a chemotopic glomerular response map. In particular, alcohols with similar chain lengths induce partially overlapping activity patterns, suggesting that not only chemically related odorants evoke similar activity patterns, but also that chemical features are not encoded by a single glomerulus but rather by a glomerular ensemble. To probe whether this coding logic applies also to the American cockroach, we tested the neural representation of a series of aliphatic alcohols with increasing chain length. A correlation analysis performed on the evoked response maps revealed that alcohols with similar length generated more similar glomerular response patterns (**Fig. 4F**). In addition, superimposing glomerular activity from different animals onto a reference antennal lobe revealed a shift in the response maps from a ventro-medial to a more dorso-lateral region of the antennal lobe upon stimulation with short-to long-chain alcohols (**Fig. 4G**). Thus, there is a topological mapping of the chemical structure (here: chain length) onto the AL geometry. These findings are consistent with previous observations in holometabolous insects such as the honeybee or the fruit fly, and indicate that, despite the divergent evolutive history, the logic behind olfactory coding inferred from holometabolous insects may also apply to (at least some) hemimetabolous ones.

### Encoding of spatial olfactory information

The American cockroach is endowed with extremely long segmented antennae, such that different antennal segments are sufficiently apart to experience different olfactory environments. Having access to receptors along the full length of a flagellum can, in principle, provide several types of information: (1) the movement of a plume towards or away from the animal could be sensed; (2) a concentration gradient along the antenna could be detected – a relevant feature since cockroaches live in shelters, where air turbulences are small, and concentrations gradients may be important; (3) the spatial structure of an odour plume could be analysed, *i*.*e*. whether the plume is spatially uniform (indicating a single odorant source), or whether the plume is spatially complex (indication distinct odorant sources). The long antennae make the cockroach an ideal model for investigating if and how the spatial structure of an olfactory stimulus could be encoded. One hypothesis would be that there is antennotopic mapping, *i*.*e*. that the brain maintains the information of proximal and distal olfactory receptors separate. In this case, we would expect a spatial segregation of proximal and distal innervation, and a mechanism to maintain that separation to higher order brain centres. An alternative hypothesis would be that spatial information is extracted through active sensing, that is by sweeping antennae in space, and proprioceptive analysis of antennal position. In this case, we would expect the highest sensitivity of the antenna to be in the distal segments, which become the most important segments for spatial analysis. A recent report from Nishino *et al*. revealed that the cockroach macroglomerulus (MG) is innervated by a group of projection neurons with an ordered subglomerular morphology (Nishino et al., 2018). By means of single cell electrophysiology, they showed that each MG projection neuron responds to a restricted antennal region. Here, by monitoring the activity across all MG uPNs at once, we tested the consequences of this finding. Our observations are consistent with the model of a functionally antennotopical glomerular organisation, and show that distal stimuli induced stronger responses in the medial portion of the macroglomerulus (*i*.*e*. close to the entry site of uPNs into the MG), whereas proximal stimulations resulted in a lateral/cortical MG activation. This finding complements the previous report (Nishino et al., 2018) showing that glomeruli are not the smallest unit of olfactory processing, but that sub-glomerular structures may convey information about stimulus location along the antennae. In addition, we observed that distal stimuli have a tendency to elicit weaker responses than proximal stimuli. Even if not statistically significant, this observation is coherent with previous electrophysiological observations, which showed that uPNs responsive to distal pheromone stimuli respond with about 60% (Hosl 1990) or 75% (Nishino 2018) of the intensity of uPNs responsive to proximal stimuli. This effect is likely due to a decrease in sensillar density along the cockroach antenna, thus to a smaller population of responsive neurons (Hösl, 1990; Watanabe et al., 2018).

Can the spatial arrangements of other common odorants be encoded by a similar mechanism in other (“ordinary”) glomeruli? Our study shows that the sensitivity to common odorants – differently from the sensitivity to periplanone B – decreases drastically along the antenna. With calcium imaging analysis, we could detect a clear response to a plume of odorant localized a few millimetres from the head, but almost no activity when the odour plume was presented to a more distal antennal section. This effect can be explained only partially by the decrease in sensillar density along the antenna from proximal to distal (Hösl, 1990; Watanabe et al., 2018). Indeed, olfactory sensilla density may explain up to 2-fold decrease in sensitivity along the antenna, but not the 5-to-10-fold signal decrease that we observed (**Fig. 5**). Such a low sensitivity in the distal portion of the antenna suggests that spatial analysis along the antenna might be more important for the cockroach than proprioceptive analysis of a moving antenna. However, this does not exclude that both mechanisms may be used in parallel, or in different ecological contexts. In our experiments, antennae were immobilized in order to control stimulus location, and to avoid movement artefacts. It may be that proprioceptive analysis has a modulatory effect on antennal lobe activity, *i*.*e*. that distal stimulation would elicit stronger responses in the antennal lobe if (and only if) the antennae were moving.

Most odour molecules interact with multiple ORN types with different affinities, activating multiple glomeruli across the antennal lobe. However, differently from the MG, ordinary glomeruli are not equipped with multiple uPNs with stereotyped sub-glomerular morphology, such that all cognate OSNs terminate onto a single uPN. Thus, if the spatial arrangement of an odorant plume could be perceived by the cockroach olfactory system, it is unlikely that it relies on the same mechanism expressed by the MG, because each single uPNs per glomerulus could not encode stimulus location of common odorants. Glomerular subcompartments (and thus antennotopic position of the stimulus) may, however, be encoded by other AL neurons such as local interneurons or multiglomerular PNs. Indeed, each glomerulus is innervated by multiple mPNs, which do not always innervate the whole glomerulus, but rather ramify in restricted sub-glomerular portions (Malun et al., 1993). This type of architecture is reminiscent of the spatial coding arrangement observed for uPNs in the MG (Nishino et al., 2018). In the MG, a large-receptive field uPN (L1) covers the entire MG volume, while multiple small-receptive field uPNs cover limited portions of the MG. In the rest of the antennal lobe, each glomerulus is fully innervated by one uPN (in analogy with the L1-PN system for the MG), while multiple mPNs innervate several glomeruli, although only partially (in analogy with the small-receptive field PNs found in the MG, but spatially distributed across multiple glomeruli). In order to test this hypothesis, future studies of mPN single cell labelling and electrophysiology are required. If, indeed, spatial analysis were performed by mPNs, this would create an interesting dichotomy in odour coding, with a segregation of information across different tracts: space would be coded in mPNs and travel along the lateral ALTs, while odour quality would be coded across uPNs, and travel within the medial ALTs. Similar parallel systems are present across insects and sensory modalities (Galizia and Rössler, 2010).

### Spatial and temporal signal integration

Two odorants, A and B, originating from the same source are likely to travel within the same odour plume (Nowotny et al., 2013; Stierle et al., 2013). If such a homogeneous plume crosses the antenna, the same odorant mixture will be experienced by the whole antenna in the same way, triggering the activation of the receptors for A and for B at the same annuli, in synchrony. Hence, the information of the activated OSN population will reach the AL with the same latency producing a glomerular response map for the synchronous odour object AB. Conversely, if odours A and B originate from different sources, they will result in an asynchronous plume, where components A and B do not travel in synchrony, and reach the antenna at different locations and at different times. Thus, the experience of an asynchronous mixture will evoke the activity of two different OSN ensembles in different spatial or/and temporal locations (one for odorant A and one for odorant B), which will reach the AL with different latencies, depending on the spatio-temporal separation of the two sources. In this second case, a dynamic response map would arise, with a glomerular code shifting from the most proximal component, *e*.*g*. A, to the most distal one, *e*.*g*. B. In our analysis, we measured that a stimulus in a certain portion of the antenna induced a response in the AL PNs, whose latency is a function of the axonal length of the activated OSNs, with a travelling speed of ∼0.37 m/s. Thus, a stimulus separation of 40 mm along the antennal surface would produce a delay greater than 100 ms in uPNs response time. Estimates obtained from moth PNs show that multiple signals can be temporally integrated when separated by less than 60-80 ms (Tabuchi et al., 2013). This means that two stimuli far apart on the antenna may not be subject to temporal integration, thus supporting the hypothesis that a long antenna would allow for instantaneous discrimination of synchronous versus asynchronous mixtures (Nowotny et al., 2013; Stierle et al., 2013).

### Conclusive remarks

In conclusion, we show that glomeruli are complex functional units with a subglomerular structure, and suggest that this arrangement may contribute to the ability to encode spatial information of olfactory stimuli. While we confirmed that space along the antenna is maintained as space within the macroglomerulus, further studies are required to fully understand if and how information provided by the antennotopic arrangement of non-pheromone receptors is maintained within the brain. We show that odour quality, odour concentration, and chemical similarity follow a combinatorial logic comparable to what is found in holometabolous insects and in mammals. Finally, proximal antennal segments induce stronger signals in the antennal lobe, whereas the distal half of the antenna evokes very weak calcium responses, raising new questions about how the cockroach uses its characteristic long antennae in olfactory coding, and active sensing. Further studies, including calcium imaging in multiglomerular projection neurons, will help us elucidating olfactory coding across species.

## Acknowledgement

This work was funded by the DFG Centre of Excellence 2117 “Centre for the Advanced Study of Collective Behaviour” (ID: 422037984). Thanks to Mark Willis for the periplanone samples. Thanks to Stephanie Neupert for experimental control software and to Silvio Widmer for assistance in the volumetric analysis of AL glomeruli.

## Supplementary Information

**Figure S1.**
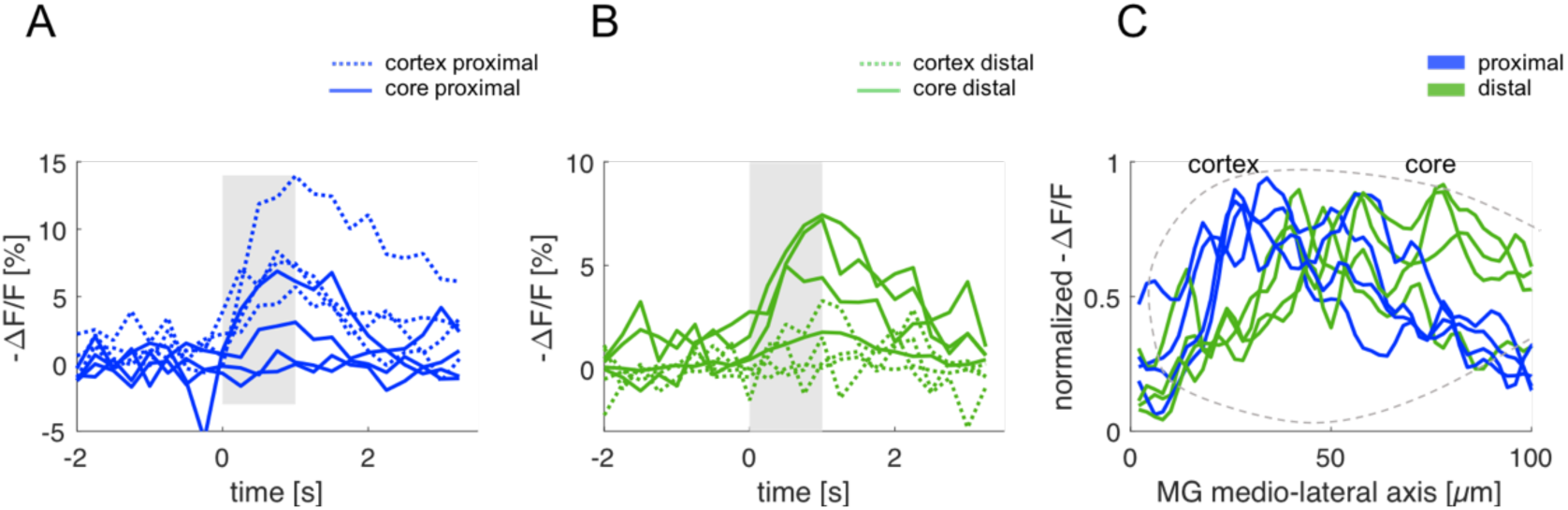
Antennotopic odour response in the macroglomerulus. Individual traces from four animals showing evoked responses in the MG core (continuous line) and cortex (dotted line) to a proximal (**A**) and a distal (**B**) stimulation. (**C**) Individual traces showing the evoked responses along the longitudinal MG axis for proximal and distal stimulation, where position 0 µm corresponds to the most lateral side and position 100 µm to the most medial side of the MG.

## Supplementary Movie Legends

**Movie 1. Bilateral antennal lobe dye injection reveals projection neurons tracts**. Image size: 1190 × 1190 µm; z-interval between consecutive frames: 2 µm.

**Movie 2. Z-stack of antennal lobe uniglomerular projection neurons labelling**. Image size: 643 × 643 µm; z-interval between consecutive frames: 3 µm.

**Movie 3. Antennal lobe spontaneous and stimulus-evoked activity**. The video shows 20 s of calcium imaging in the projection neurons of the antennal lobe. While intense activity can be observed during stimulation (green rectangle), ongoing activity can be observed also before and after stimulation. Video was acquired at 25 Hz but presented at 12.5 Hz (every second section was omitted for reduce file size); stimulus (1-octanol 10^−2^) was delivered two times for 0.1 s, at *t* = 4 s and *t* = 14 s. Colormap of raw data presented as Δ*R*_340/380_. No filtering or post-processing was applied to this video. Scale bar, 100 µm.

