## Supplemental material for "Coding of odour and space in the hemimetabolous insect *Periplaneta americana*"

#### Supplementary Information

### Figure S1

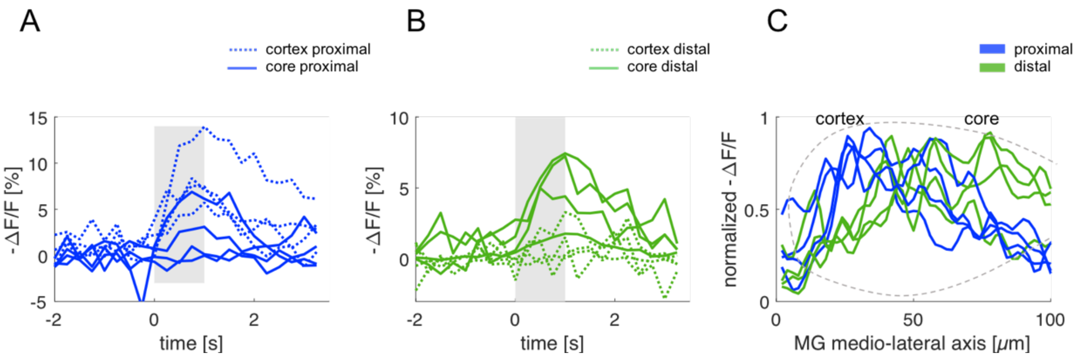


**Figure S1. Antennotopic odour response in the macroglomerulus.** Individual traces from four animals showing evoked responses in the MG core (continuous line) and cortex (dotted line) to a proximal (**A**) and a distal (**B**) stimulation. (**C**) Individual traces showing the evoked responses along the longitudinal MG axis for proximal and distal stimulation, where position 0 μm corresponds to the most lateral side and position 100 μm to the most medial side of the MG.

#### Supplementary Movie Legends

**Movie 1. Bilateral antennal lobe dye injection reveals projection neurons tracts.** Image size: 1190 × 1190 µm; z-interval between consecutive frames: 2 µm.

**Movie 2. Z-stack of antennal lobe uniglomerular projection neurons labelling.** Image size: 643 × 643 µm; z-interval between consecutive frames: 3 µm.

**Movie 3. Antennal lobe spontaneous and stimulus-evoked activity.** The video shows 20 s of calcium imaging in the projection neurons of the antennal lobe. While intense activity can be observed during stimulation (green rectangle), ongoing activity can be observed also before and after stimulation. Video was acquired at 25 Hz but presented at 12.5 Hz (every second section was omitted for reduce file size); stimulus (1-octanol $10^{-2}$) was delivered two times for 0.1 s, at $t = 4$ s and $t = 14$ s. Colormap of raw data presented as $\Delta R_{340/380}$. No filtering or post-processing was applied to this video. Scale bar, 100 µm.
